# High-throughput imaging of *Caenorhabditis elegans* aging using collective activity monitoring

**DOI:** 10.1101/2021.10.18.464905

**Authors:** Anthony D Fouad, Matthew A Churgin, Julia Hayden, Joyce Xu, Jeong-Inn Park, Alice Liu, Christopher Teng, Hongjing Sun, Mateo Parrado, Peter Bowlin, Miguel De La Torre, Timothy A. Crombie, Christine A. Sedore, Anna L. Coleman-Hulbert, Erik Johnson, Patrick Philips, Erik C. Andersen, Christopher Fang-Yen

**Affiliations:** Department of Bioengineering, School of Engineering and Applied Science, University of Pennsylvania, Philadelphia PA; Department of Molecular Biosciences, Northwestern University, Evanston, IL; Department of Biology, University of Oregon, Eugene, OR

## Abstract

The genetic manipulability and short lifespan of *C. elegans* make it an important model for aging research. Widely applied methods for measurements of worm aging based on manual observation are labor intensive and low-throughput. Here, we describe the Worm Collective Activity Monitoring Platform (WormCamp), a system for assaying aging in *C. elegans* by monitoring activity of populations of worms in standard 24-well plates. We show that metrics based on the rate of decline in collective activity can be used to estimate the average lifespan and locomotor healthspan in the population. Using the WormCamp, we assay a panel of highly divergent natural isolates of *C. elegans* and show that both lifespan and locomotor healthspan display substantial heritability. To facilitate analysis of large numbers of worms, we developed a robotic imaging system capable of simultaneous automated monitoring of activity, lifespan, and locomotor healthspan in up to 2,304 populations containing a total of ~90,000 animals. We applied the automated system to conduct a large-scale RNA interference screen for genes that affect lifespan and locomotor healthspan. The WormCamp system is complementary to other current automated methods for assessing *C. elegans* aging and is well suited for efficiently screening large numbers of conditions.

## INTRODUCTION

Studies in the roundworm *C. elegans* have led to a number of important discoveries about the basic mechanisms of aging (Kenyon, 2010). A number of aging-associated pathways discovered or extensively studied in *C. elegans*, including insulin/insulin-like signaling, DAF-16/FOXO, TOR, and dietary restriction, have proven to be conserved in other animal models and possibly humans (Friedman and Johnson, 1988; Kenyon et al., 1993; Fraser et al., 2000; Lee et al., 2003; Hansen et al., 2005; Olsen et al., 2006; Uno and Nishida, 2016; Mack et al., 2018).

Traditional assays of aging in *C. elegans* require direct visual observation of worms and manual manipulation with a wire pick to assess lifespan (Lee et al., 2003). Manual lifespan assays are low-throughput and lack potentially important information about behavior and/or health during aging. These limitations of manual assays create barriers for large-scale efforts to discover novel genetic, pharmacological, or environmental interventions to slow or mitigate the effects of aging.

To address these limitations, automated methods have been described for automatically assessing *C. elegans* aging. These methods include image analysis of worms cultured as individuals on separate substrates, as a population on a standard agar plate, or in microfluidic devices that may be used to observe worms individually or in populations (Stroustrup et al., 2013; Xian et al., 2013; Zhang et al., 2016; Churgin et al., 2017; Pittman et al., 2017; Saberi-Bosari et al., 2018; Pitt et al., 2019; Rahman et al., 2020).

Our laboratory previously developed a microfabricated multi-well device called the WorMotel in which worms are observed in arrays of up to 240 wells, each holding one animal (Churgin et al., 2017). Data from the WorMotel was used to analyze the trajectories of behavioral decline during aging in wild-type and mutant strains (Churgin et al., 2017). Although the WorMotel method excels at providing detailed longitudinal information, it is difficult to scale to tens of thousands of animals due to the need to manually fabricate the substrates and prepare individual worms and wells. As a result, it is not ideally suited for large-scale genetic and/or pharmacological screens.

To address these limitations, we describe an experimental method for assaying *C. elegans* aging by monitoring populations of worms on standard 24-well microplates (**Figure 1, Supplemental Figure 1.1, Video 1**). We define metrics that, based on the collective behavioral activity from each well, can be used to predict the mean lifespan and locomotor healthspan (the time during which the locomotor activity of an individual exceeds a certain threshold) of each population. We then developed an improved automated imaging system capable of serially monitoring many plates at a time. We show that this assay, the Worm Collective Activity Monitoring Platform (WormCamp), enables recordings of up to 2,304 populations simultaneously and therefore facilitates measurements and screens for *C. elegans* lifespan and locomotor healthspan.

**Figure 1:**
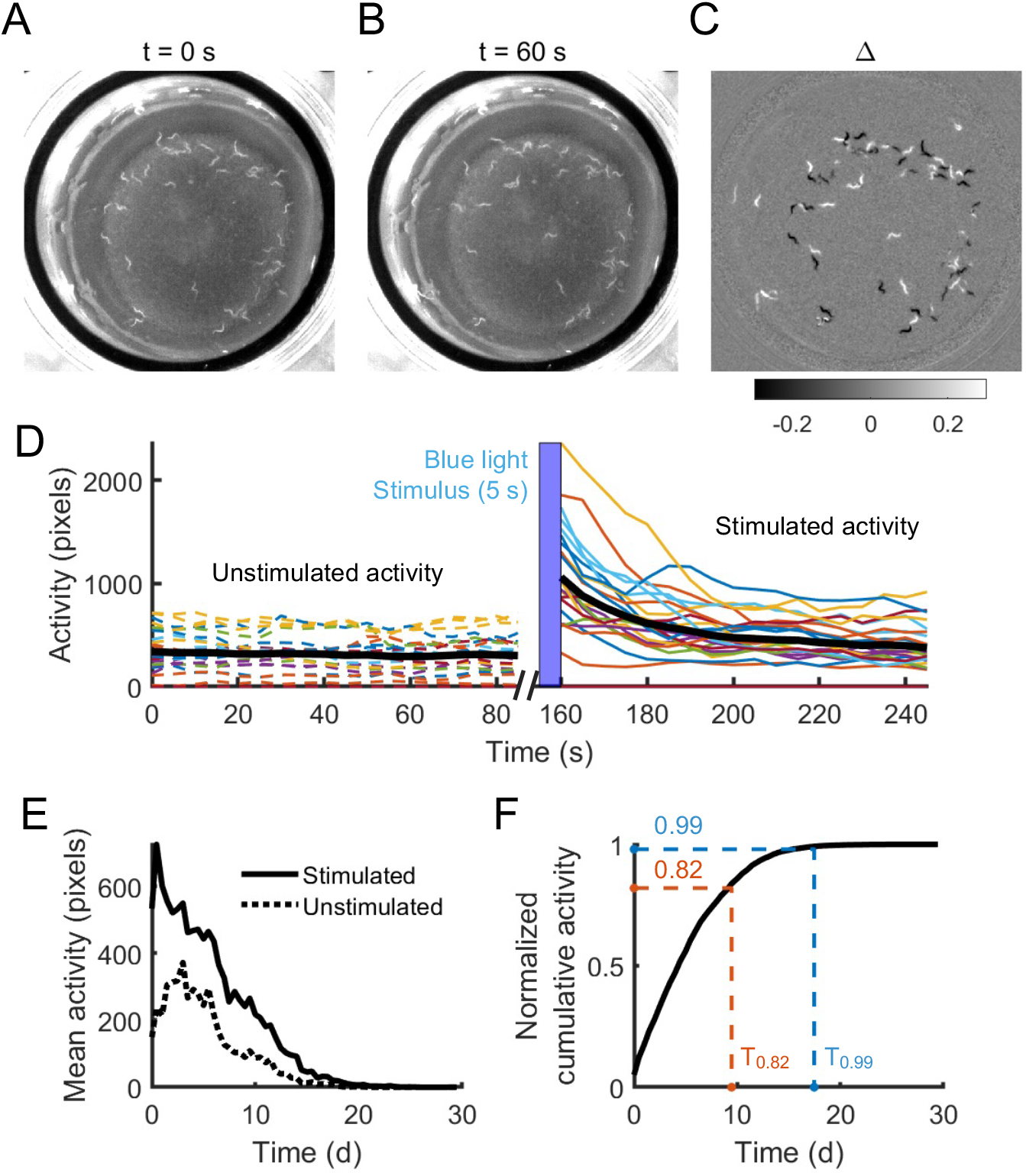
A 24-Well plate-based method for measuring *C. elegans* population lifespan and healthspan. (A) A population of approximately 30 worms in a single well from a 24-well plate. (B) The same well 60 s later. (C) The difference image between the two frames. Gray represents pixels with no change; black represents pixels that decreased in intensity and white represents pixels that increased in intensity. Pixels that increased or decreased in intensity by more than 15% were summed separately, and the activity between two frames was defined as the lesser of the two sums. (D) Quantification of activity of 24 worm populations during a 5 min recording in which images were recorded every 5 seconds (similar to Churgin et al, 2017). Image pairs 1 min apart were analyzed as in (A-C). A blue LED stimulation occurred at t = 155 s; activity before this time is referred to as unstimulated, activity after this time is stimulated. Data is censored between t = 90 s and t = 155 s as the difference image would be computed from one stimulated and one unstimulated frame, and between t = 245 s and t = 300 s because no paired image exists to subtract. (E) Activity curves for a full 30 day experiment for one population. The stimulated and unstimulated activity at any timepoint in an activity curve is the mean of the corresponding activity that occurred during the 5-minute session. (F) Normalized cumulative activity corresponding to the stimulated activity in (E). Times to reach NCA = 0.82 and NCA = 0.99 are defined as the healthspan and lifespan estimates, respectively.

## RESULTS

### Aging phenotypes can be inferred from collective activity

In the previously described WorMotel (Churgin et al., 2017), each well contains a single animal, allowing longitudinal analysis of behavior and lifespan. By contrast, we measure the collective activity of the population in each well in the WormCamp. At any time, the collective activity is equal to the number of surviving animals multiplied by the average individual activity. We asked how the collective activity from worm populations in the WormCamp might be used to assess metrics of aging, including lifespan and locomotor healthspan.

To understand the relationship between collective activity and lifespan or locomotor healthspan, we first simulated collective activity by summing individual worm activity data previously acquired using the WorMotel (Churgin et al., 2017). We used data from 8 independently prepared 240-well plates containing a variety of conditions, including wild type animals, long lived mutants such as *daf-2* (Kenyon et al., 1993), *age-1* (Friedman and Johnson, 1988), and *tax-4* (Apfeld and Kenyon, 1999), and short lived mutants such as *daf-16* (**Figure 2**).

**Figure 2:**
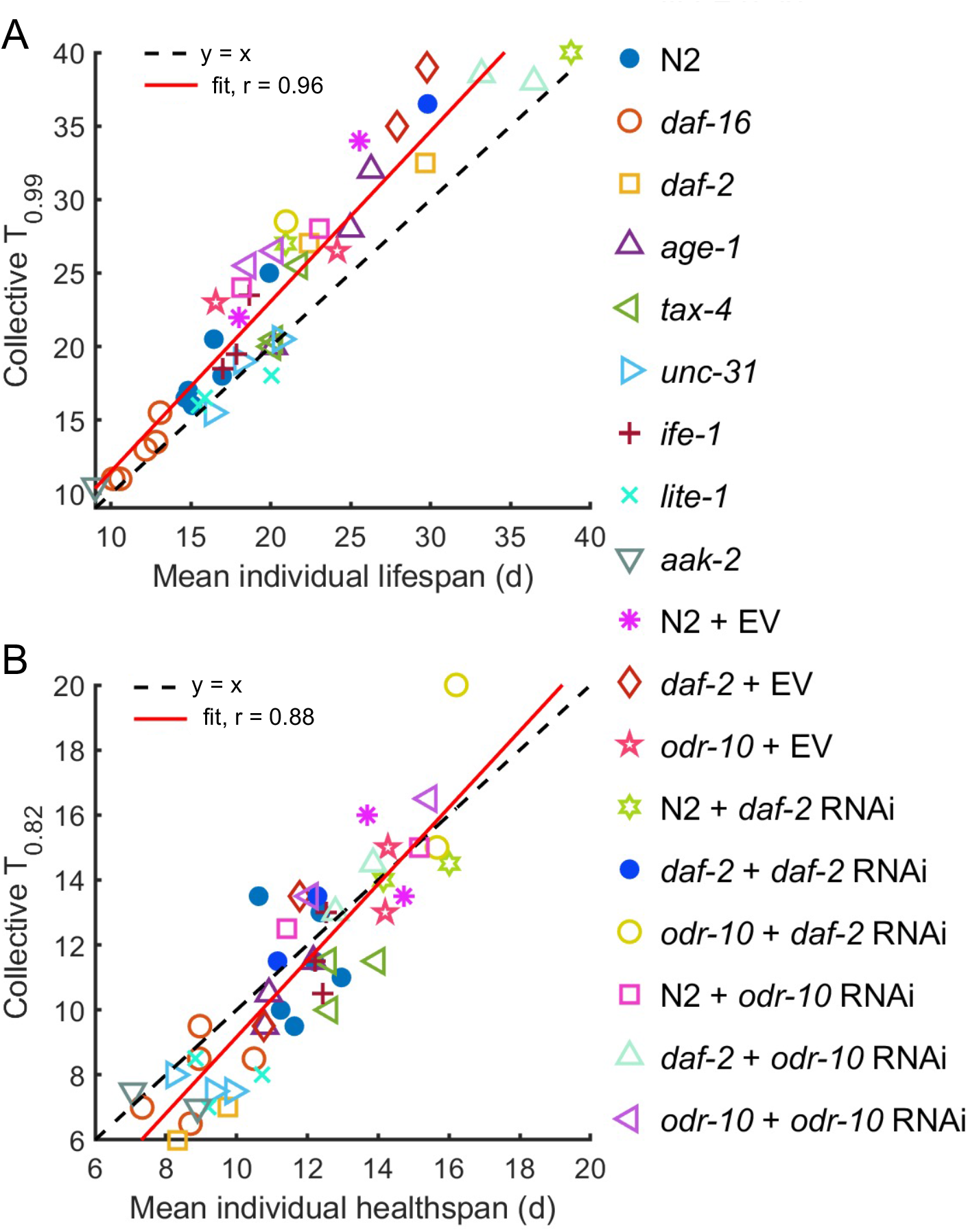
Simulations of collective activity curves from individual worm activity. (A) Correlation between the optimized lifespan estimator (T_0.99_) and true mean lifespan of a simulated populations of 30 worms in WorMotels (Churgin et al, 2017). Each population contains worms of one genotype. (B) Correlation between the optimized healthspan estimator (T_0.82_) and the true mean healthspan.

We summed up to 30 individual worm activity measurements to obtain the stimulated collective activity A(t) of the population for each recording session, analogous to that measured from a population of worms cultured together, as in a WormCamp (Figure 1B, C). We then calculated the normalized cumulative activity (NCA) for each population over time, defined as

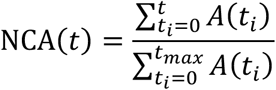

where the t_i_ are the time points at which activity was measured and t_max_ is the time of the final measurement, which exceeds the maximum lifespan in the population. Note that NCA(0) = 0 and NCA(t_max_) = 1, *i.e*., the cumulative activity monotonically rises from 0 to 1. Next, we defined a metric T_×_ = NCA^-^(x) where NCA^-1^ is the inverse function of NCA and x is a threshold between 0 and 1. The metric T_×_ corresponds to the time at which the normalized cumulative activity equals x (**Figure 1F**).

We asked whether the metric T_×_ can be used to approximate true lifespan of a population. We calculated the correlation and root mean square error (RMSE) between T_×_ and true mean lifespan for a range of values of x from 0 to 1. We found that x=0.99 gave the maximum best linear fit correlation coefficient (r = 0.96) and x = 0.97 gave the minimum RMSE (**Supplemental Figure 2.2B, D**). These results show that our T_0.99_ metric based on collective activity is a good estimator of lifespan.

Next, we asked whether the metric can also be used to estimate locomotor healthspan. Although healthspan in general lacks a consensus definition (Kaeberlein, 2018), one way to quantify locomotory health of individual worms is to determine the total time that an animal’s activity exceeds a threshold. This healthspan definition has also been referred to as the total days healthy (TDH) (Jushaj et al., 2020). In this work, we define the locomotor healthspan of an individual worm as the time during which the animal’s activity exceeded one-third of its 90^th^ percentile activity level (**Supplemental Figure 2.1**). Conceptually, this value represents the time during which the worm was at least moderately active.

As we did for lifespan, we chose x to minimize RMSE between T_×_ and mean locomotor healthspan, resulting in a value of x = 0.82. The correlation coefficient between the two variables was also near its maximum value of about r = 0.88 near x = 0.82. We therefore selected T_0.82_ as our healthspan estimator (**Supplemental Figure 2.2A, C**).

Having validated the concept of collective activity based on simulated data, we next performed experiments with actual *C. elegans* populations in single wells of 24-well plates. We plotted the mean activity value for each well before and after the blue light stimulus (used to evoke activity) for each imaging period. As expected, collective activity decreased during aging (**Figures 1C, D; 3A, B**).

**Figure 3:**
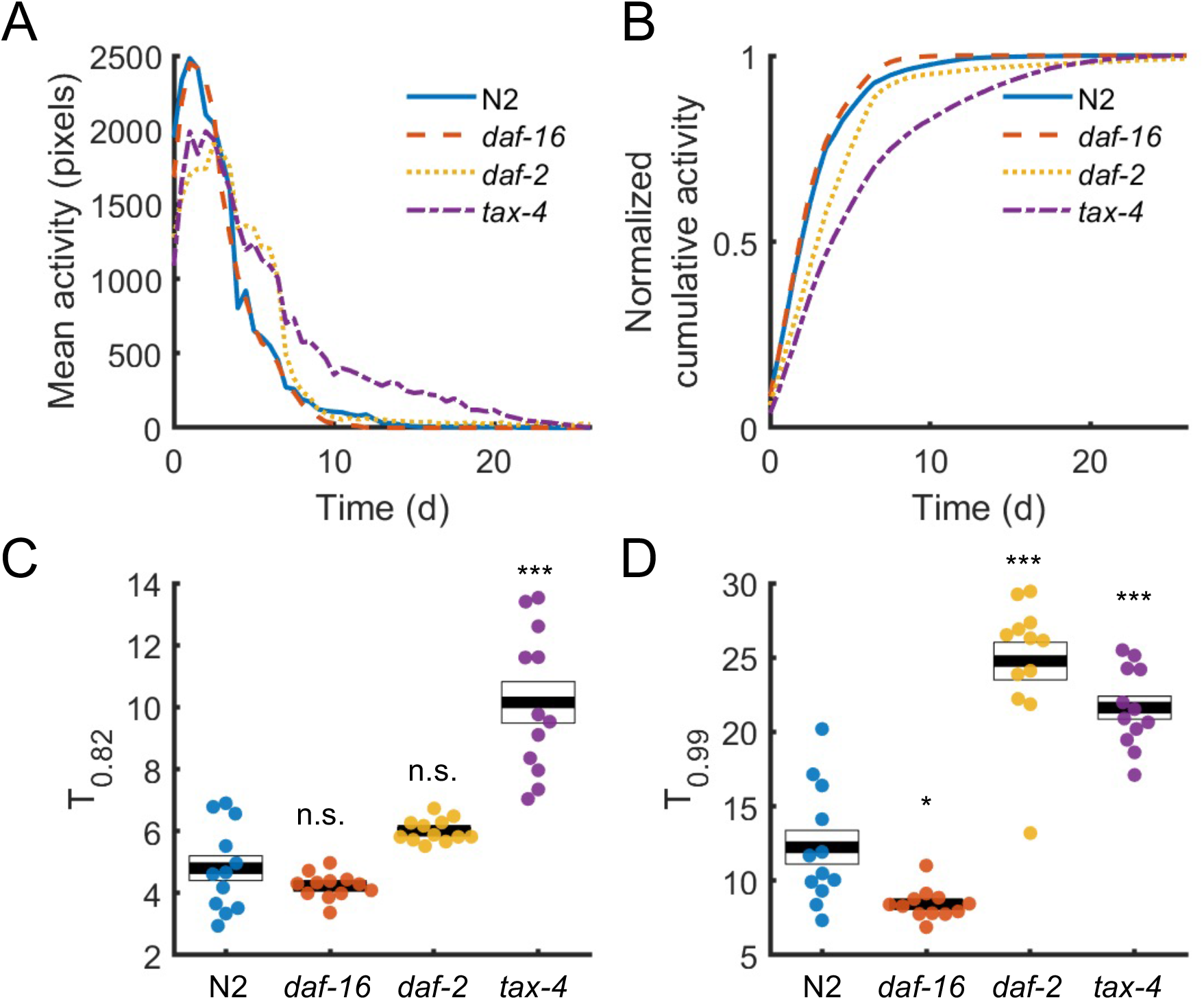
WormCamp assessment of mutants with known aging phenotypes. Mean activity (A) and normalized cumulative activity curves (B) for each of the four genotypes tested. Healthspan estimates (T_0.82_) and lifespan estimates (T_0.99_) for all wells prepared are shown in C and D, respectively. * p < 0.05, ** p < 0.01, *** p<0.001 vs N2, one-way ANOVA followed by Bonferroni-corrected post-hoc comparisons. Black bars are the group mean, white boxes indicate group SEM. 12 wells (populations) were prepared for each condition; two plates with a total of 48 wells were prepared in total.

We sought to verify that the WormCamp provides expected results for well known aging mutants. We performed 24-well plate experiments with wild-type animals, short-lived *daf-16* mutants, and long-lived *tax-4* and *daf-2* mutants. We found that we could robustly detect differences in mean population lifespan and locomotor healthspan between these strains. Lifespan and locomotor healthspan inferred from collective activity measurements were in good agreement with that from simulating population data from individual worm data (**Figure 2A-B, 3B-C**).

These results demonstrate that a metric defined by the normalized cumulative activity can be used to accurately infer average lifespan and locomotor healthspan in populations.

### A robotic imaging system enables high-throughput screening using the WormCamp

To increase experimental throughput, we sought to develop a method for imaging many plates concurrently. Various robotic imaging systems are potentially suitable for this task, including plate handling robots that move plates to a stationary imaging system (Churgin et al., 2017), and those that use motorized stages to move a camera to image stationary plates (Alisch et al., 2018; Pitt et al., 2019). We selected the moving-camera approach to reduce the rate of mechanical faults we had previously experienced with plate-handling robots.

Our system, which we call WormWatcher, is a customizable integrated hardware and software solution for monitoring experiments concurrently. We constructed two systems containing 3-dimensional motorized stages and platforms, one with a capacity of 77 plates and another with a capacity of 96 plates. Mounted to each stage is an assembly containing a camera, three high-powered blue stimulation LEDs mounted to a heat sink, and an internally mirrored acrylic box which promotes uniformity of blue light illumination on the plate. Red LED strips are mounted to the base of the mirror box to provide dark field illumination (**Figure 4**). Although this illumination method creates some spatial nonuniformity in dark field illumination across the plate (**Appendix I – Figure 2A-C**), we found that a well’s mean gray intensity had negligible influence on either the lifespan or healthspan estimators (r^2^ ≈ 0.02 for either; **Appendix I – Figure 2D-F**). As part of a study of the genetic basis of aging (Luyten et al., 2016), we used the robotic imaging systems and the WormCamp method to screen the clones in the Ahringer RNAi library (Source Bioscience)(Kamath et al., 2003) for their effects on lifespan and locomotor healthspan (Appendix I).

**Figure 4:**
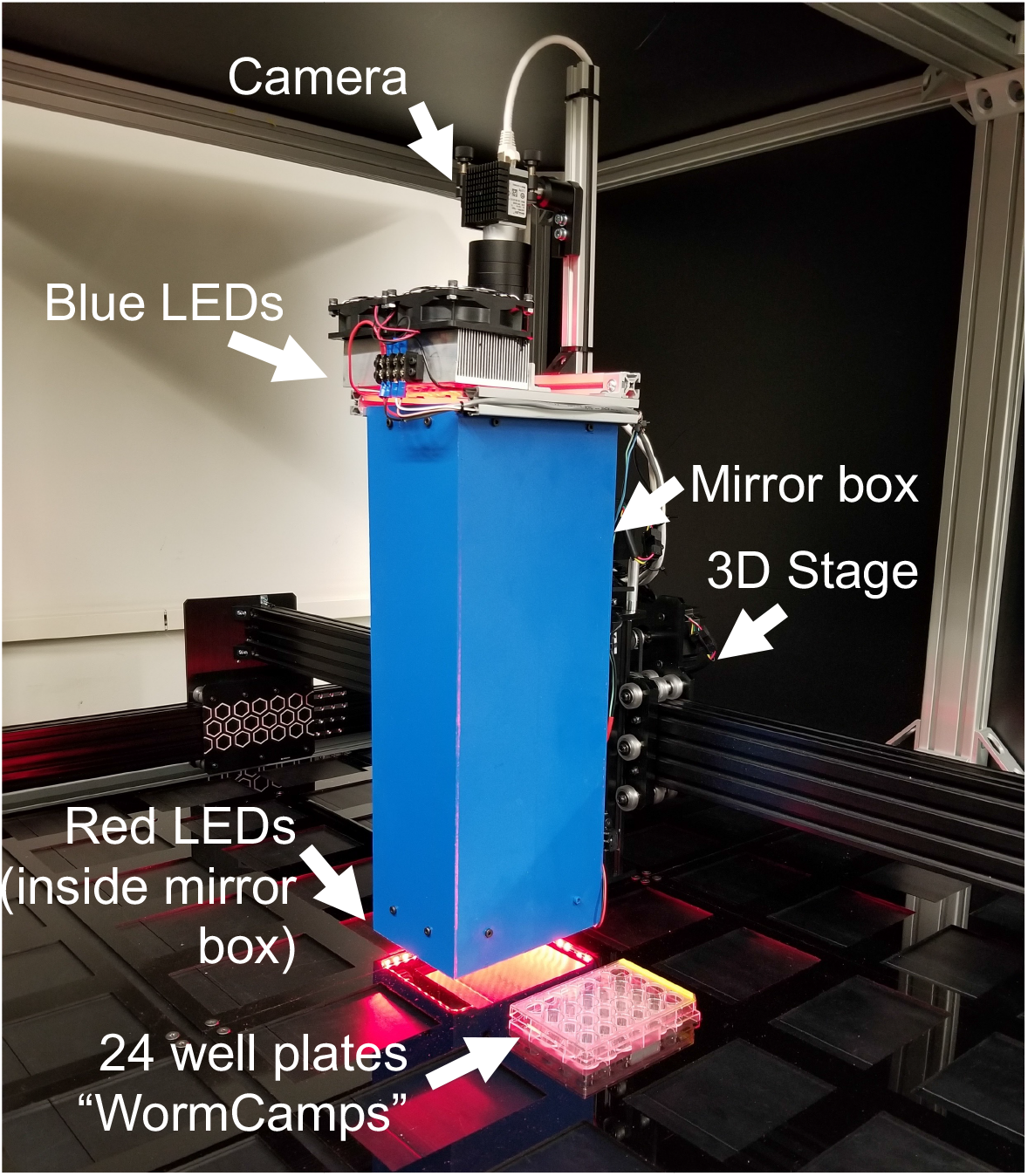
Robotic imaging systems. Image of a WormWatcher robot representative of the two robots used in our screen. Components of the imaging apparatus are labeled.

### WormCamp measurements measure heritability of lifespan and locomotor healthspan in C. elegans wild isolates

As part of a study of natural variation in aging, we applied the WormCamp method and our WormWatcher robots to assay lifespan and locomotor healthspan in a representative panel of 12 wild *C. elegans* strains. These 12 strains are highly diverged from one another and from the laboratory reference strain, together capturing 74.4% of the total known variation in *C. elegans (Cook et al., 2017; Lee et al., 2021)*.

We assayed 253 populations containing a total of approximately 7,500 animals. We estimated that the broad-sense heritabilities of lifespan and locomotor healthspan are 0.17 and 0.31, respectively (**Figure 5A, B**).

**Figure 5:**
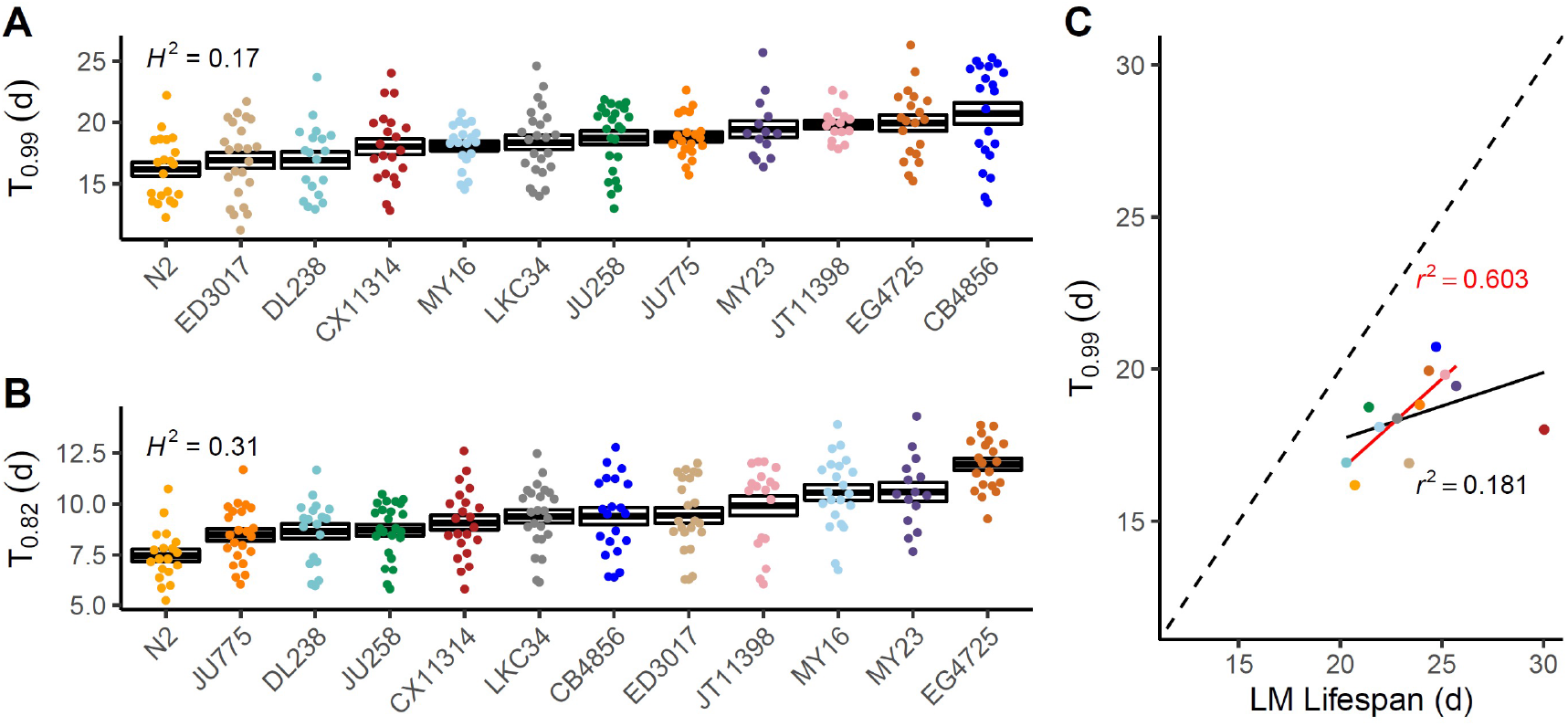
Comparison of WormCamp and Lifespan Machine assessments of wild isolate lifespan. (A) lifespan (T_0.99_) and (B) healthspan estimates (T_0.82_) for 12 wild isolates. (A, B) Points represent individual well trait estimates and are colored by strain, boxes correspond to the first and third quartiles, and the black bars represent median trait values. Low quality wells are censored and outlier values within a strain are removed by Tukey’s fences (n = 17 - 26 wells per strain). (C) A correlation between WormCamp (T_0.99_) and Lifespan Machine lifespan assessment. Points are median lifespan estimates from replicate wells (WormCamp) or plates (Lifespan Machine) and are colored by strain. Spearman’s rho for all strains (black) and without CX11314 (red)

Previous reports have described large variability in lifespan measurements between different laboratories and between different assay techniques (Lucanic et al., 2017). We asked how WormCamp measurements of the 12 wild strains compare with those acquired by another method. To that end, we performed experiments on the same 12 strains using the Lifespan Machine, an automated population-based assay of lifespan using flatbed scanners (Stroustrup et al., 2013) (**Figure 5**). We found a high degree of correlation (Spearman correlation coefficient 0.78) between the measurements acquired in two different labs despite using vastly different experimental assays. These results increase confidence that the differences in aging we observe in wild isolates are robust and reproducible between laboratories.

### Optimization of well preparation and automated assessment of experiment quality

Our large-scale screen presented the challenge of ensuring high data quality from thousands of experiments containing tens of thousands of images. Manual inspection of portions of the data indicated that the most common data aberration was the tendency of some of the worms to “escape”, *i.e*., disappear from the visible portion of the well, within approximately the first week (**Appendix I - Table 1**). Worms were later found desiccated on the vertical walls of the wells. Other problems that affected smaller portions of the dataset were bacterial or fungal contamination and the presence of progeny due to failure of floxuridine to sterilize the animals.

Because disappearance substantially affected the overall quality of this dataset, we developed an automated method for estimating the number of worms in each well so that we could exclude them from further analysis.

First, we attempted to develop a simple method based on gray level or adaptive thresholding that could reliably differentiate worms from background. However, these efforts were unsuccessful, primarily due to variations in the texture of the bacteria lawn and lighting intensity and uniformity across each plate.

Recent studies have shown that machine learning and convolutional neural network (CNN) based approaches can reliably identify worms in variable quality images (Hakim et al., 2018; Bornhorst et al., 2019; Bates et al., 2021). We trained a CNN to differentiate worms from background in 20×20 pixel kernels extracted from our images (see above). The 20×20 classification kernel is scanned across an image of a well in about 0.1s, determining whether each location tested is more likely to be worm or background (**Figure 6A-C**) and reports the worm fraction (WF), the fraction of tested locations deemed likely to contain worms. This WF is highly correlated with the number of worms counted manually in an image (**Figure 6D**).

**Figure 6:**
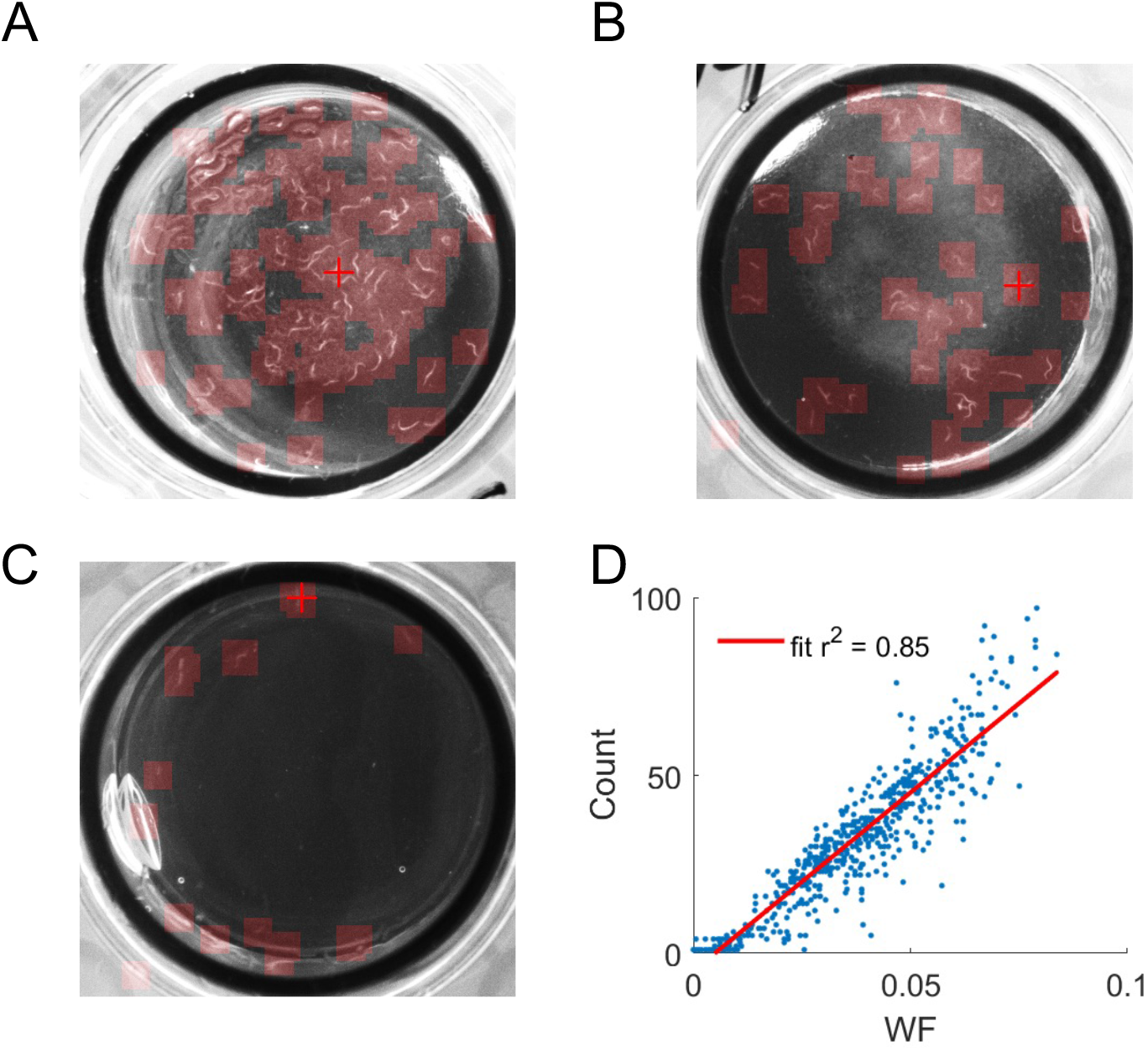
CNN-based worm detection and counting. (A-C) Demonstrations of CNNbased worm detection on crowded (A), ideally populated (B), and empty (C) WormCamp wells. Red overlay denotes image regions that were classified as worm. (D) Correlation between worm fraction (WF) and the number of worms labeled manually in test images that were not used for training.

Application of the CNN to our entire dataset revealed that WF generally decreases over time (**Appendix I – Figure 4A-C**). We expected a decrease over each 30-day experiment because many worms appeared to disintegrate after death, precluding their detection by the CNN (**Video 1**). Nonetheless, large decreases in worm count over the first 10 days of imaging were unexpected and a common problem, with most wells experiencing a decrease by day 10, and some wells experiencing a loss of most of the worms (**Appendix I – Figure 4B**).

We next sought to determine the cause of worms escaping. We found that the percentage of worms remaining at day 10 was not meaningfully correlated with the worm density (initial WF) or with the lighting intensity of the well (**Appendix I – Figure 4D-E**). We next asked whether wells prepared with the bacteria lawn touching the edge of the well were associated with increased escaping, as has been observed on standard NGM plates (Stiernagle, 2006). To address this question, we prepared WormCamp plates in which some wells contained a strictly centered lawn (**Figure 7A**), some wells contained a lawn that was centered but touching the edge (**Figure 7B**), and some wells contained a lawn that was off center and substantially touching the edge (**Figure 7C**). Populations in the latter two well types experienced significantly more escaping than populations placed on strictly centered lawns (**Figure 7D-E**). These results show that ensuring that the bacteria lawn does not touch the edge of a well is important for reducing escaping and producing accurate activity measurements.

**Figure 7:**
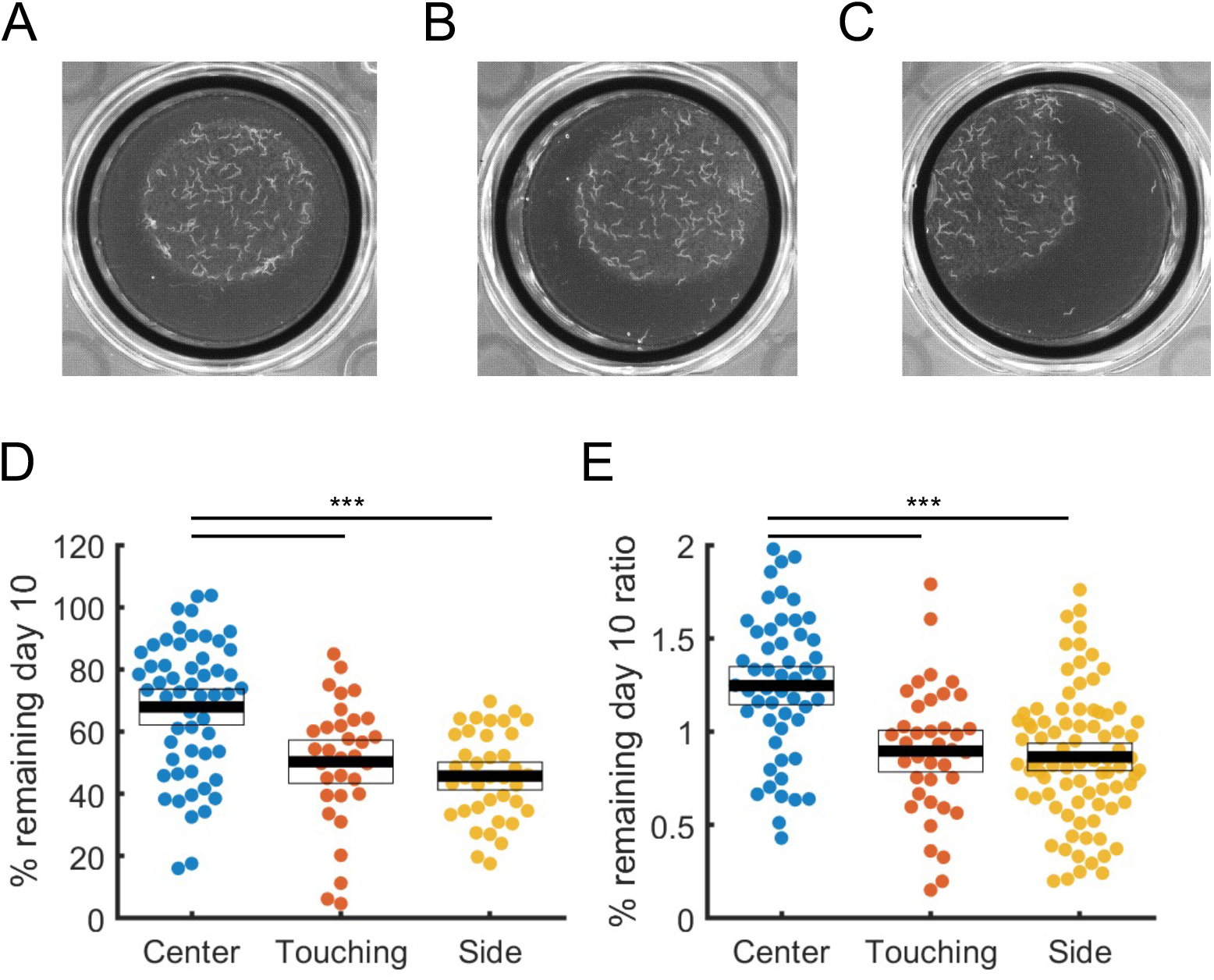
Centering the bacteria lawn within the well reduces rate of worms escaping from wells. (A-C) Images of WormCamp wells prepared with the bacteria lawn centered without touching the edge of the well (A), somewhat off-center, slightly touching the edge (B), very off-center and substantially touching the edge (C). (D) percentage of WF retained at day 10 for worms cultured on each lawn type. (E) Ratio of the mean percentage retained for the plate (E). For D and E, 10 plates (240 wells) were prepared in total, each containing some wells with each lawn type. N = 58, 33, and 91 wells of each type, respectively. The remaining wells were excluded due to contamination and/or progeny. *** p<0.001, one-way ANOVA followed by Bonferroni-corrected post-hoc comparisons.

## Discussion

Our WormCamp method is an efficient way to conduct high-throughput measurements of *C. elegans* aging. Our technique consists of (1) an experimental procedure based on standard 24-well plates for measuring aging parameters in *C. elegans* populations and (2) an automated imaging system for serially monitoring plates containing thousands of populations. By validating aging metrics using ground truth data from individual worm aging trajectories acquired using the WorMotel and matching those results to actual data from worm populations grown in 24-well plates, we show that key aging parameters can be extracted from collective population activity data. We have also shown that our automated imaging system can be used to assay natural variation in aging among wild isolate strains, and to screen for genes with effects on lifespan and healthspan.

The WormCamp method has several important advantages compared to the WorMotel method (Churgin et al 2017). First, compared to a 240-well WorMotel, the WormCamp has a several times larger number of worms per plate (960 worms if N=40 per well), and can therefore test more conditions at a given population size per comparison. Another advantage is that no PDMS device fabrication or plasma cleaning are necessary for the WormCamp because commercial multiwell plates are used. Agar, food bacteria, and worm loading procedures are also simpler due to the smaller number of wells. Finally, less manual editing is required for data analysis.

The WormCamp method also has disadvantages compared to the WorMotel. Since worms are not monitored individually in the WormCamp, there is no longitudinal tracking of behavior and no direct measurements of activity, lifespan, or healthspan. Although we have defined accurate estimates for lifespan and healthspan based on a wide variety of mutant strains, we cannot rule out the possibility that there exist strains in which the relationship between cumulative activity and lifespan or healthspan as described here do not hold.

Given these relative strengths and weaknesses, the WormCamp and WorMotel are best viewed as complementary methods. The WorMotel is well suited for acquiring high-resolution longitudinal data for aging and behavior, including more accurate measures of lifespan and healthspan, and the WormCamp is well suited for efficiently screening a large number of strains or conditions.

A number of other methods for automated assays of aging have been described. These include microfluidic devices (Xian et al., 2013; Saberi-Bosari et al., 2018; Rahman et al., 2020), in which worms are imaged in a liquid environment. Among the advantages of these methods are that worms can be imaged at high resolution and without the presence of the chemical sterilization. However, microfluidics expose worms to an atypical environment and depend on specialized fluid handling equipment that can be difficult to scale to the levels we have achieved using the WormCamp.

Methods for automated aging studies of worms on solid substrates include the scanner-based Lifespan Machine (Stroustrup et al., 2013) and the WormBot (Pitt et al., 2019). These methods both assay lifespan by tracking individual worms in periodically acquired images. The Lifespan Machine acquires images using modified flatbed scanners, but the WormBot moves a camera to each of a number of six-well plates using a threedimensional stage similar to our robotic imaging system. These methods are most accurate when worms are spaced out in larger plates, and when worms are near death and move very little between imaging sessions. Compared to these methods, the

WormCamp has the advantage of increased throughput due to the confinement of worm populations in relatively small wells, and the disadvantage of not providing direct measures of individual worm fates as discussed above.

We used the WormCamp to assess aging phenotypes of 12 representative wild isolates of *C. elegans*. We found that mean lifespan and locomotor healthspan traits have a broad-sense heritability of 0.32-0.34 (Fig. 7), a value greater than estimates of heritability of lifespan in humans (Herskind et al., 1996). Future work will use the automated robotic system to extend our measurements to hundreds of wild strains to identify new genetic loci that modulate aging across natural populations.

We used the WormCamp to perform a large-scale screen for RNAi knockdowns affecting aging. Previous RNAi screens for lifespan genes have been reported (Lee et al., 2003; Hamilton et al., 2005; Hansen et al., 2005). However, the manual lifespan assays used to conduct these screens are limited by manual labor and unable to identify genes influencing differences in health during aging. Our method is capable of identifying new modulators of aging and distinguishing effects on lifespan from those on healthspan. The ability to efficiently measure both lifespan and healthspan will enable additional genetic and pharmacological analyses of aging.

## METHODS

### Strains

The following strains were used in this study: N2, CF1038: *daf-16(mu86) I*, CB1370: *daf-2(e1370) III*, TJ1052: *age-1(hx546) II*, PR678: *tax-4(p678) III*, DA509: *unc-31(e928) IV*, KG1180: *lite-1(ce314) X*, RB754: *aak-2(ok524) X*, CX32: *odr-10(ky32) X*, Wild isolates: CX11314, MY16, LKC34, JU775, DL238, ED3017, JU258, MY23, JT11398, EG4725, CB4856. All strains were maintained at 15-20°C using standard methods (Sulston 1983). All experiments were carried out at approximately 20°C unless otherwise stated.

Experiments with the WorMotel platform were performed as described (Churgin et al., 2017). Experiments on the Lifespan Machine were performed according to published protocols (Stroustrup et al., 2013; Mark et al., 2021).

### Preparation of 24-well plates

Media for 24-well WormCamp plates is prepared identically to standard NGM plates (Stiernagle 2006) with agar replaced by 25 g/L gellan gum (Gelrite, Sigma Aldrich), which creates a gel with greater optical transparency than agar does. We included 200 μM 5-fluoro-2’-deoxyuridine (floxuridine, Sigma Aldrich) in the media to prevent growth of progeny. For non-RNAi WormCamp experiments, 200 ng/mL streptomycin was added to the media. For RNAi WormCamp experiments, 1 mM IPTG and 200 μg/mL carbenicillin were included in the media. 1 mL of media was added to each well using a 25 mL electronic pipette. To provide hydration, the region between the microplate wells was filled with approximately 30 mL of a solution of 20 g/L gellan gum and 3 g/L NaCl.

After media solidified, 50 μL of a DA837 or HT115 overnight bacteria suspension was added to each well for non-RNAi and RNAi experiments, respectively.

Worms were synchronized by hypochlorite bleaching and added to each well approximately 40 hours after plating (Stiernagle 2006). L4-stage worms were suspended in NGM and approximately 30 worms were added to each well using a P1000 manual pipette. Although some wells may contain more or less than 30 worms, we verified that the initial worm count was not correlated with aging outcomes (**Appendix I – Figure 4D**). After allowing excess liquid to dry, we sealed plates using Parafilm laboratory film.

### Static Rig Image Acquisition

For static (non-robotic) systems, images were captured with an Imaging Source DMK 72AUC02 camera (2592 × 1944 pixels) equipped with a lens (Fujinon HF25SA-1, 2/3” 25mm focal length, f/1.4 C-Mount, Fujifilm Corp., Japan). We used Phenocapture imaging software or MATLAB (MathWorks) to acquire time lapse images through a USB connection. Static imaging rig experiments were carried out under dark-field illumination using four 4.7” red LED strips (Oznium, Pagosa Springs, CO) positioned approximately 2” below the plate (Churgin et al. 2017; Churgin and Fang-Yen 2015). For static imaging experiments, plates were imaged at a rate of 0.2 Hz for 30-minute imaging periods initiated every 12 h, and a 10 s blue light stimulus occurred at approximately 15 min.

### Blue LED Stimulation

Blue light stimulation was carried out as previously described (Churgin et al. 2017). Briefly, two or three high-power blue LEDs (Luminus Phlatlight PT-121, Sunnyvale, CA) were mounted to an aluminum heat sink for static or automated serial monitoring experiments, respectively. Blue LEDs were secured atop a mirrored acrylic box which served to generate uniform illumination to the plate. A solid-state relay (6325AXXMDS-DC3, Schneider Electric, France) controlled by MATLAB through a NIDaq USB-6001 I/O device (National Instruments, Austin, TX) was used to drive the LEDs at a current of 18 A using a DC power supply (29950 PS, MPJA Inc.).

### Image Processing

Images were processed in MATLAB in a manner similar to that previously described (Churgin et al. 2017). Briefly, images acquired 60 seconds apart were subtracted. For each region of interest (well), pixels for which the gray-scale intensity value either increased or decreased by more than 15% were counted separately. We expect worm movements to create both increased and decreased pixels (Figure 1C), whereas artifacts such as a temporary increase or decrease in lighting intensity create only one type of changed pixel. Therefore, we used the lesser of the sums of increased and decreased pixels as the activity between two pixels. The activity of the well for a frame was defined as the number of pixels changed within each well.

Each imaging period consisted of a set of frames acquired before or after blue light stimulation (**Figure 1D**). For each imaging period, the spontaneous and stimulated activity for each well was defined as the average activity value over all frames in that imaging period before or after blue light stimulation, respectively.

### Robotic Imaging System and software

The robotic plate imaging system is based on a commercially available computer numerical control (CNC) kit (SMW3D OX CNC). Briefly, the CNC kit was assembled according to the manufacturer’s instructions but with a custom imaging and illumination assembly attached to the stage in place of the spindle. Microcontroller-driven stepper motors move the carriage to visit each plate location twice per day. An array of laser-cut slots in an acrylic base plate define the locations of each plate. The software is optimized for the 24-well plate assay described above, but the robot is also compatible with any 128 mm x 86 mm plate, such 96-well plates, 384-well plates, or 240-well WorMotels (Churgin et al., 2017). The first robot has a footprint of 1.5 m x 1.25 m and can image an array of 11 x 7 = 77 plates containing 1848 populations. The second robot has a footprint of 1.2 m x 2.4 m and can image an array of 8 x 12 = 96 plates or 2304 populations. The 3D stage is customizable, and smaller or larger designs are possible. Users interact with the robot via software written in C++ and Python which integrates robot movements, image acquisition, blue light stimulation, plate maintenance, error checking, and data processing.

The custom imaging assembly consists of a camera and lens (see *Image Acquisition*), blue LED and heat sink (see *Blue LED Stimulation*), and acrylic mirror box (see *Blue LED Stimulation*) mounted to 24”-long 20-mm aluminum extrusion (**Figure 4**). A single 3’ red LED strip (Oznium, Pagosa Springs, CO) was mounted to the inner side of the bottom of the mirror box. The aluminum extrusion frame was mounted to the z-axis of the CNC machine.

For automated serial monitoring experiments, each plate was imaged approximately every 12 h for 5 min. During acquisition, frames were recorded every 5 s and a 5 s blue light stimulus was applied approximately 2.5 min from the start of acquisition. Images were saved and processed by a 64-bit computer with a 3.40 GHz Intel Core i7 processor and 8 GB of RAM running Windows 10. Images were analyzed using custom-written C++ and Python software.

### Worm identification CNN Training

A convolutional neural network (CNN) was developed to identify worms in dark field images. The network was designed and trained using PyTorch (Paszke et al., 2019) and accepts a 20×20 grayscale image as input. The structure consists of three convolutional layers, made up of 6, 16, and 32 filter kernels, followed by three linear layers comprised of 200, 84, and 2 nodes. Annotated data for training and testing was generated by first drawing boxes around worms in 474 images of individual WormCamp wells taken from experiment days 4, 10, or 20 containing 0 - 80 worms each. Additional annotated data was prepared using 50 images of WorMotels (Churgin et al, 2017) containing 50-240 worms each (one per well), taken from experiment days 3 or 20.

To generate training and testing datasets, annotated images were first downsampled to 2/3 of its original resolution (such that the length of a worm is approximately 20 pixels) and then 20×20 images from random locations within each annotated image were exported. Each 20×20 image was assigned to the worm category if the center of any worm annotation lay within the middle 14×14 pixels of the 20×20 image, or the background category otherwise. In total, 67,000 worm training images, 67,000 background training images, 16,000 worm test images and 16,000 background test images were generated. Test images came from different WormCamp plates or WorMotels than training data and were never used for training.

The resulting CNN was converted to Open Neural Network Exchange (ONNX) format and is used by the deep neural network (DNN) module of OpenCV in out C++ analysis code to find worms. The final test accuracy of the CNN was 0.94 (sensitivity = 0.96, specificity = 0.92, PPV = 0.93, NPV = 0.96). To decrease the prevalence of false negatives, an additional confidence threshold of 1.6 was applied to the CNN worm node’s output, resulting in an accuracy of 0.90 (sensitivity = 0.83, specificity = 0.97, PPV = 0.97, NPV = 0.85). To locate worms within a larger image, the CNN is applied to 20×20 pixel subsets of the image, classifying each as worm or background in this manner.

### Wild strain experiments

A total of 16 WormCamp plates were prepared with wild strains. Each WormCamp contained two wells of each of 11 wild strains and two wells of N2 worms. The position of each strain within each plate was varied between plates. WormCamps were then placed on the WormWatcher robots for at least 25 days and analyzed as described in the text. Application of automated quality filters removed 33% of the 386 populations studied, chiefly for reasons of escaping (17% of all populations) or insufficient initial activity (7% of all populations). See Appendix I for more information on automated quality filters.

## Acknowledgements

Some strains were provided by the CGC, funded by NIH Office of Research Infrastructure Programs (P40 OD010440). This project has received funding from the National Institutes of Health (R21-AG053638) and the European Union’s Horizon 2020 research and innovation program under grant agreement 633589.

---

**Supplemental Figure 1.1:**
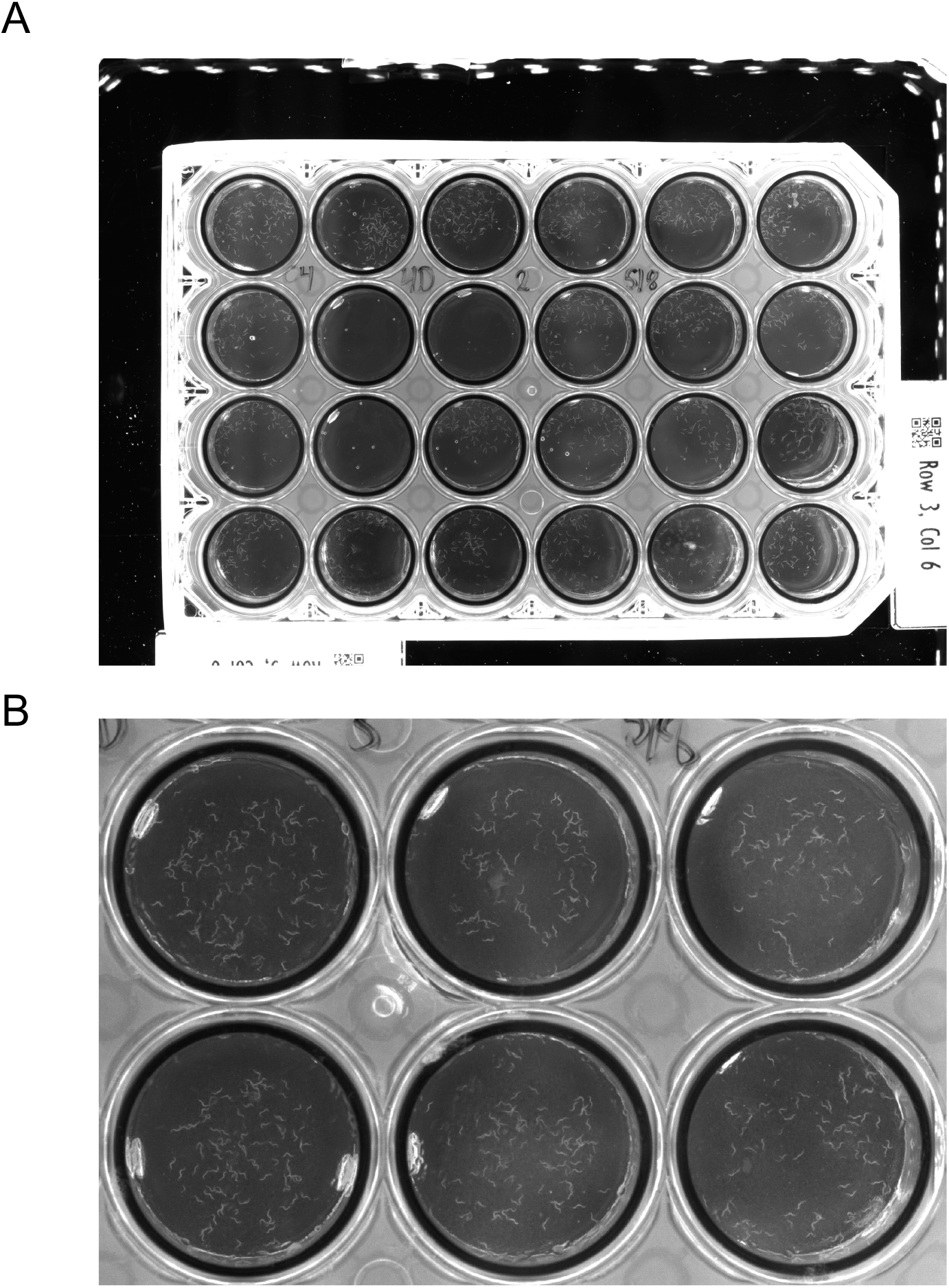
(A) Raw image of a 24-well WormCamp plate. (B) Six wells, with worms visible in all wells. See also Video 1.

**Supplemental Figure 2.1:**
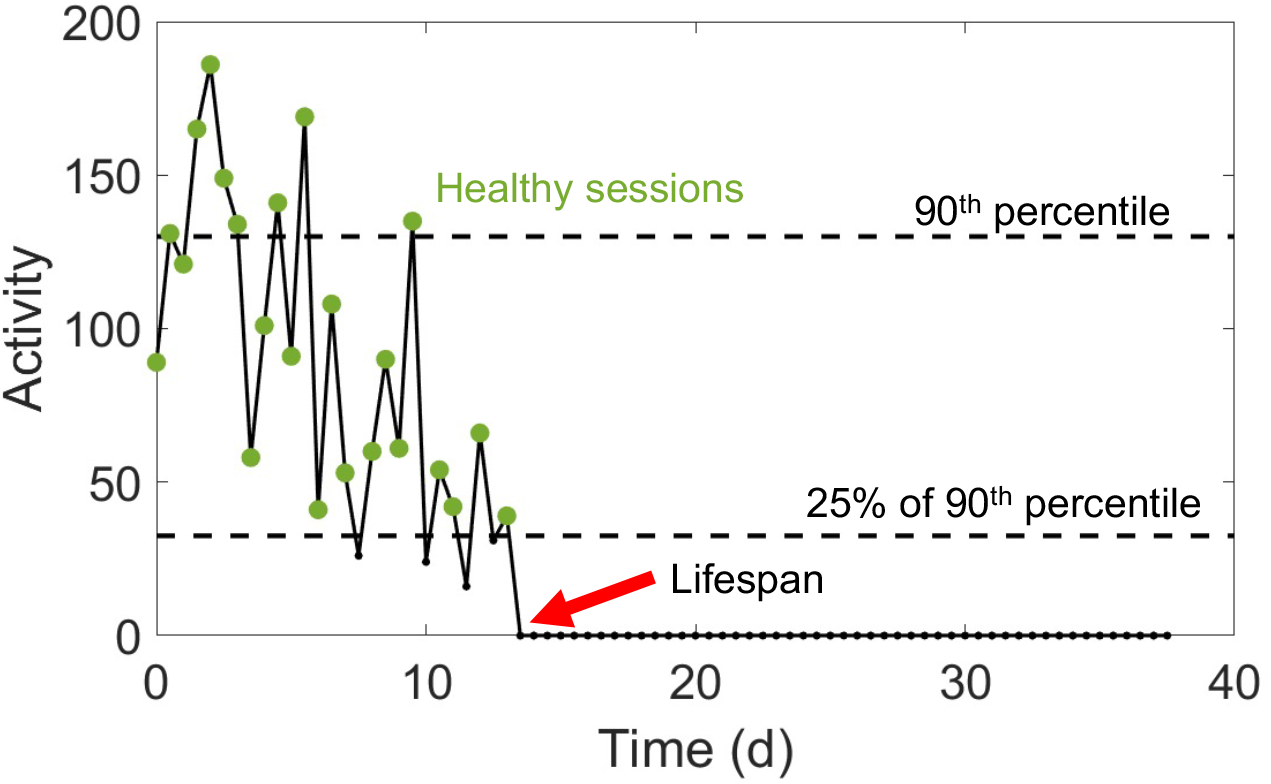
A definition of locomotor healthspan. Definition of healthspan for individual worms cultured individually in the WorMotel. Each point along an individual worm’s activity curve is the maximum stimulated activity recorded during the corresponding imaging session (Churgin et al., 2017). Healthy sessions (activity greater than 25% of the 90^th^ percentile of the activity curve) are marked with green dots. The upper dotted line indicates the 90^th^ percentile of activity, and the lower dotted line indicates the threshold for deeming a session healthy. In this example, the animal was deemed healthy on 23 sessions (healthspan = 11.5 days) and never observed to move after session 27 (arrow, lifespan = 13.5 days).

**Supplemental Figure 2.2:**
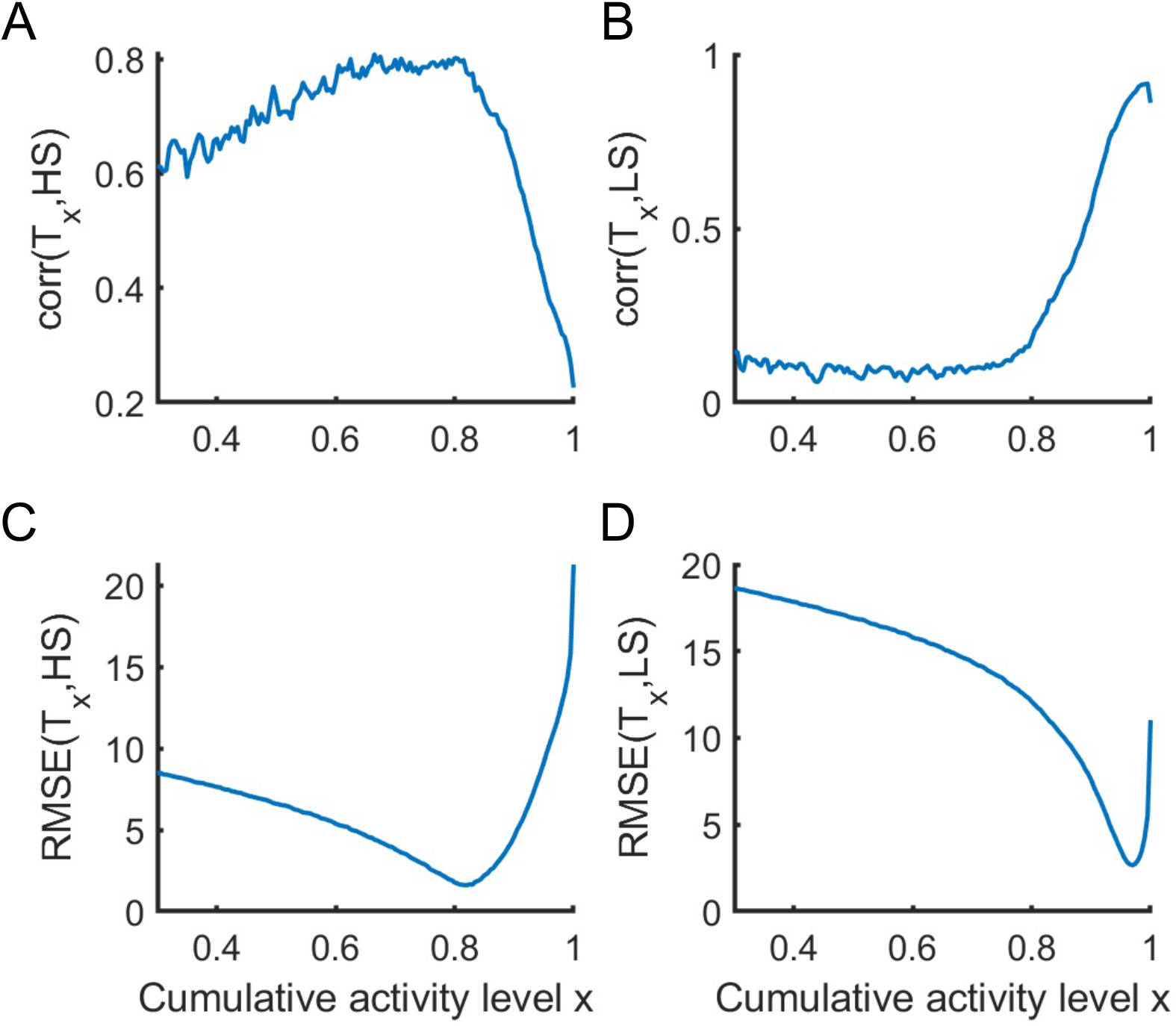
Optimization of parameters for estimation of mean lifespan and healthspan from simulated collective activity. (A) Correlation between T_×_, the time taken for the normalized cumulative activity of the simulated population to reach the xth fraction of the total, and the actual mean healthspan. (B) Correlation between T_×_ and the actual mean lifespan. Maximum: x = 0.99, correlation = 0.91. (C) Root mean squared error between T_×_ and the actual median healthspan. Minimum: x = 0.82, RMSE = 1.6 days. (D) Root mean squared error between T_×_ and the actual mean lifespan. Minimum: x = 0.97, RMSE = 2.7 days.

## APPENDIX I: An RNAi screen for aging outcomes using the WormCamp method and WormWatcher Robots

To prepare WormCamps for RNAi we used bacterial clones from the Ahringer RNAi Library (Source Bioscience, Nottingham, UK). For each 96-well plate from the library, clones on the plate were first transferred using a 96-well replicator device (V&P Scientific) to fresh 96-well plates containing LB agar (15 g/L agar and 20 g/L LB) with 200 μg/mL carbenicillin. Plates were incubated approximately 12 h at 37°C. Bacterial cultures were then transferred from the 96 well plate by adding 50 μL of LB broth with each well, suspending bacteria by pipetting, and depositing bacteria in one well of a deep 24-well plate (EK-2239, E&K Scientific, 10 mL capacity per well) along with an additional 7 mL of LB containing 200 μg/mL carbenicillin. The deep plates were covered and incubated with shaking at 37°C for approximately 12 h. To induce RNA, we added IPTG to a final concentration of 1 mM and incubated bacterial cultures for 2 h. The deep well plates were spun down on a centrifuge and most of the supernatant was removed using a vacuum aspirator. We pipetted 50 μL of the dense bacteria slurry into each well of the 24-well WormCamp plate.

Worms were added to WormCamp plates within several days of plating RNAi bacteria. To prepare animals we created synchronized worm cultures using hypochlorite bleaching, washed late L4 or young adult animals into tubes using NGM buffer, and pipetted a suspension of worms into each WormCamp well.

Each 24-well plate was prepared with four EV controls. 40% of plates also contained *daf-2* RNAi and *daf-16* RNAi positive controls. The remaining wells each contained a unique test RNAi. In our screen, daf-16 RNAi strongly decreased relative lifespan (**Appendix I - Figure 1D**) but *daf-2* was not significantly different from EV (p=0.16, not shown). It is possible that the *daf-2* RNAi in this library does not effectively knock down DAF-2 function; RNAi libraries are known to be prone to errors (Hansen et al., 2005; Jagadeesan and Hakkim, 2018).

We assayed a total of 18,072 populations, including approximately 2,996 empty vector (EV) controls and the remainder exposed to RNAi bacteria.

Summary statistics, healthspan and lifespan estimates for the RNAi and control populations are shown in **Appendix I – Figure 1.** We expected RNAi wells to show more variability in lifespan and healthspan than EV wells, since some gene knockdowns increase lifespan and some reduce lifespan. We found that the mean standard deviation of both T_0.82_ and T_0.99_ metrics was indeed significantly higher in RNAi wells compared to EV wells (**Appendix I – Figure 1A-C**).

During screening we observed and optimized some details of WormCamp preparation, particularly to reduce worms escaping from wells (**Appendix I – Table 1 and Methods**).

**Appendix I – Table 1:** Automated assessment of data quality.

| Quality issue | Assessment metric | % wells passing |
| --- | --- | --- |
| Initial setup | Initial WF > 0.02 | 86 % |
| Escaping | WF remaining at day 10 is > 50% initial | 83 % |
|  | WF remaining at day 20 is > 30% initial | 88 % |
| Progeny | WF remaining at day 10 is < 130% initial | 93 % |
|  | WF remaining at day 20 is < 130% initial | 94 % |
|  | Activity reaches 0 by end of experiment | 90 % |
| **Overall** |  | **55 %** |

**Appendix I – Figure 1:**
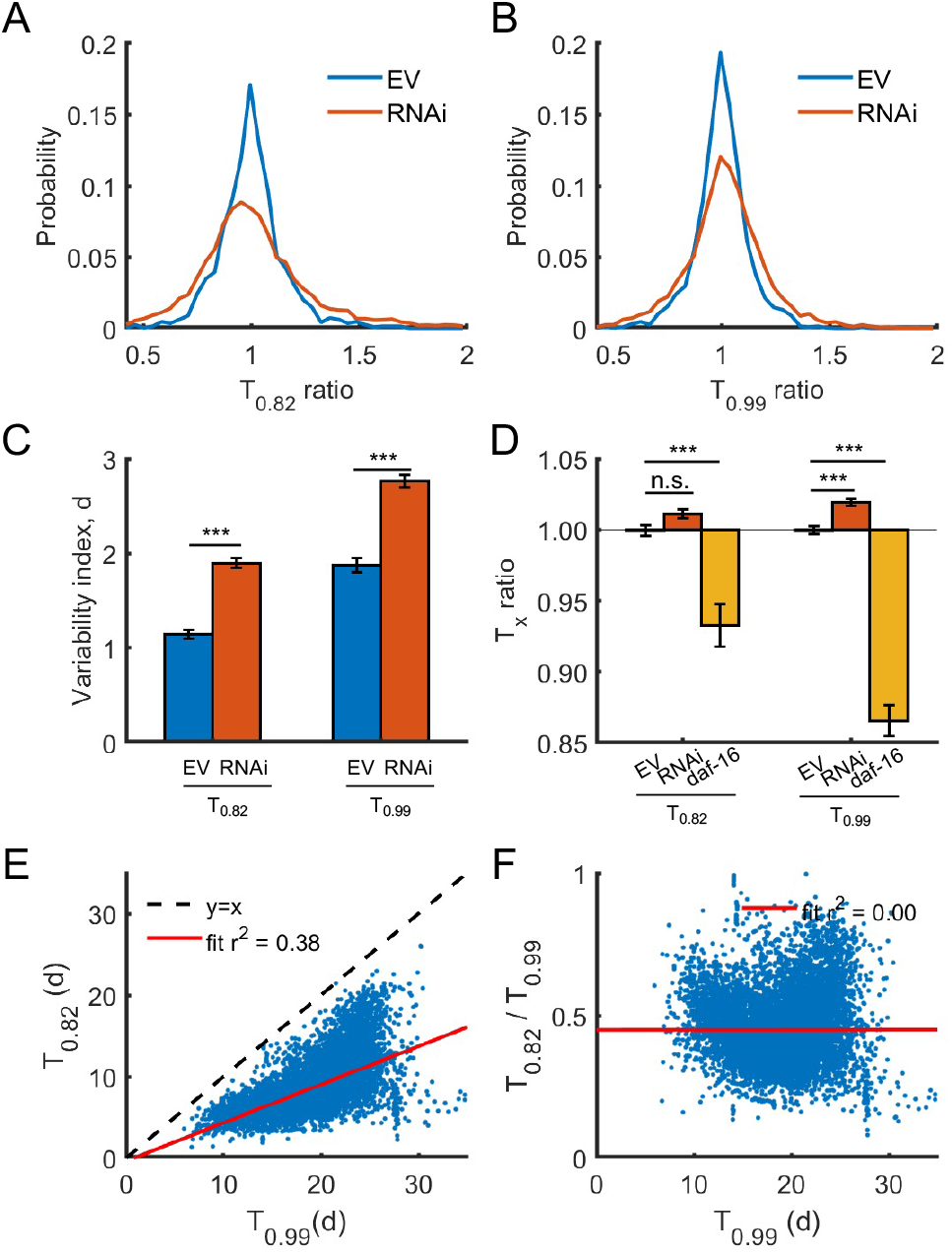
RNAi screen for ageing outcomes. (A, B) Histograms showing the distributions of T_0.82_ (A) and T_0.99_ (B) ratios across all RNAis and EV controls included in the screen. Both metrics are normalized to the mean of the four EVs on each plate. Wells were excluded if they did not pass quality tests (see text), or more than two of the four EVs were excluded due to poor quality. (C) Variability index, the mean plate-wise standard deviation of the T_0.82_ and T_0.99_ distributions, is significantly higher for RNAi wells than EV wells (*** p < 0.001, two-sample t-test). (D) Mean T_0.82_ and T99 ratios for EV wells, RNAi wells, or *daf-16* RNAi (positive control) wells from the screen (*** p<0.001, one-way ANOVA followed by Bonferroni-corrected post-hoc comparisons). (E) Correlation between lifespan and healthspan estimates for each well. (F) Correlation between lifespan estimates and the healthspan ratio T_0.82_/T_0.99_.

**Appendix I – Figure 2:**
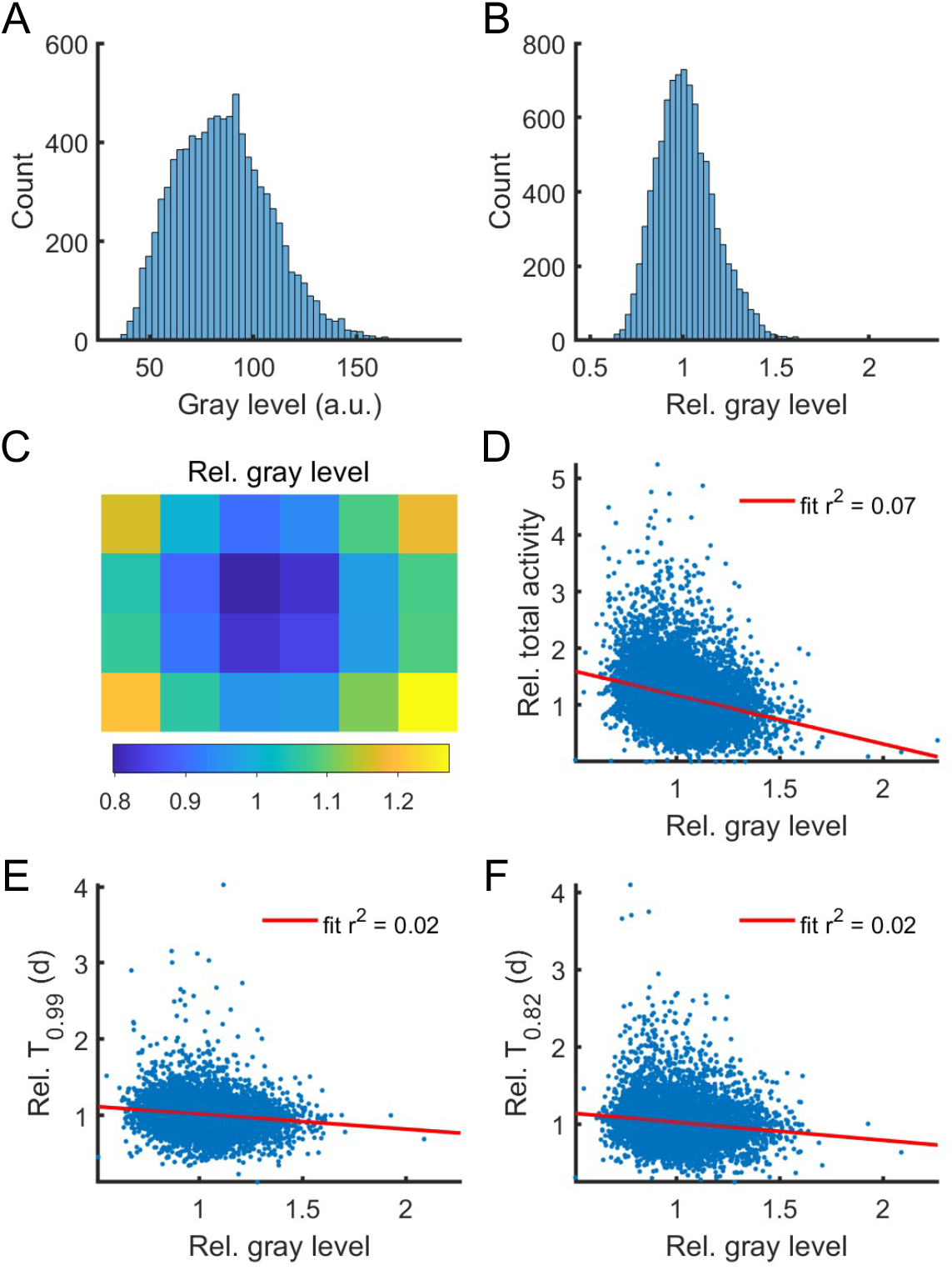
Nonuniform lighting intensity does not affect key experimental measurements. (A, B) Histograms showing the distribution in mean well grayscale intensity (A) and intensity relative to the plate average (B) of all wells in the screen. (C) Plate-relative intensity averaged across all wells in the same position on the 24-well (4 row x 6 column) plate. Note that intensity is lowest in the center of the plate, which is also furthest from the red LED lights mounted to the moving mirror box. (D-F) Correlation between relative intensity and relative total activity, T_0.99_ and T_0.82_. For all fits, p < 0.001 and r^2^ < 0.08.

**Appendix I – Figure 3:**
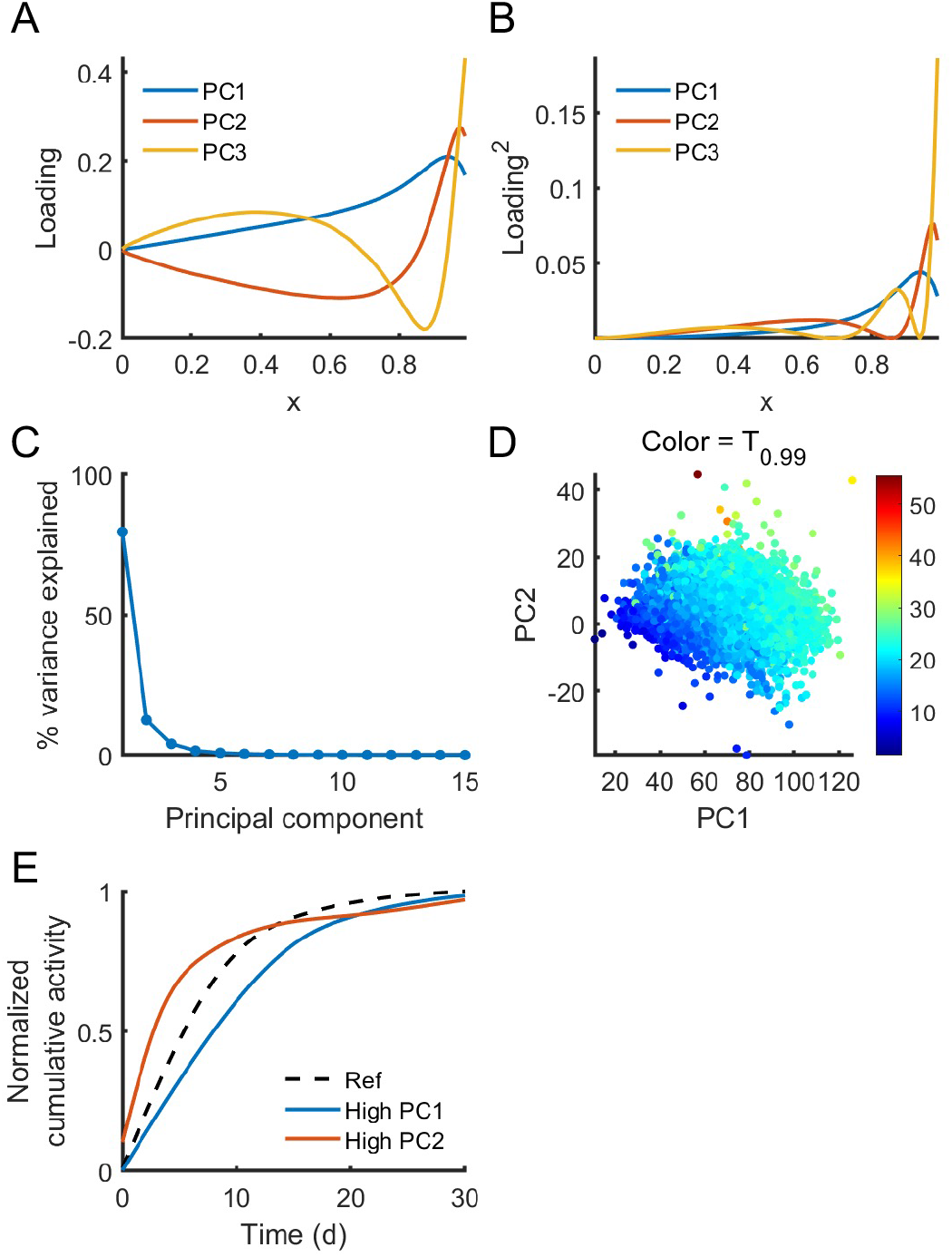
Principal component analysis of screen dataset. PCA was performed using all possible estimators T_×_ as independent variables. (A, B) Loading and square of the loading of the first three principal components. (C) Percentage of variance explained by principal components 1 through 15. (D) PC1 value vs PC2 value of each well, with the color map indicating the T_0.99_ of that well. Note higher T_0.99_ is associated with both higher PC1 and higher PC2, as expected from (A). (E) Hypothetical normalized cumulative activity curves for a reference well, a well higher in PC1, which would tend to arise from increases in all T_×_ according to (A), and a well high in PC2, which would arise from a well with increased T_×_ for x > 0.85 but decreased T_×_ otherwise.

**Appendix I – Figure 4:**
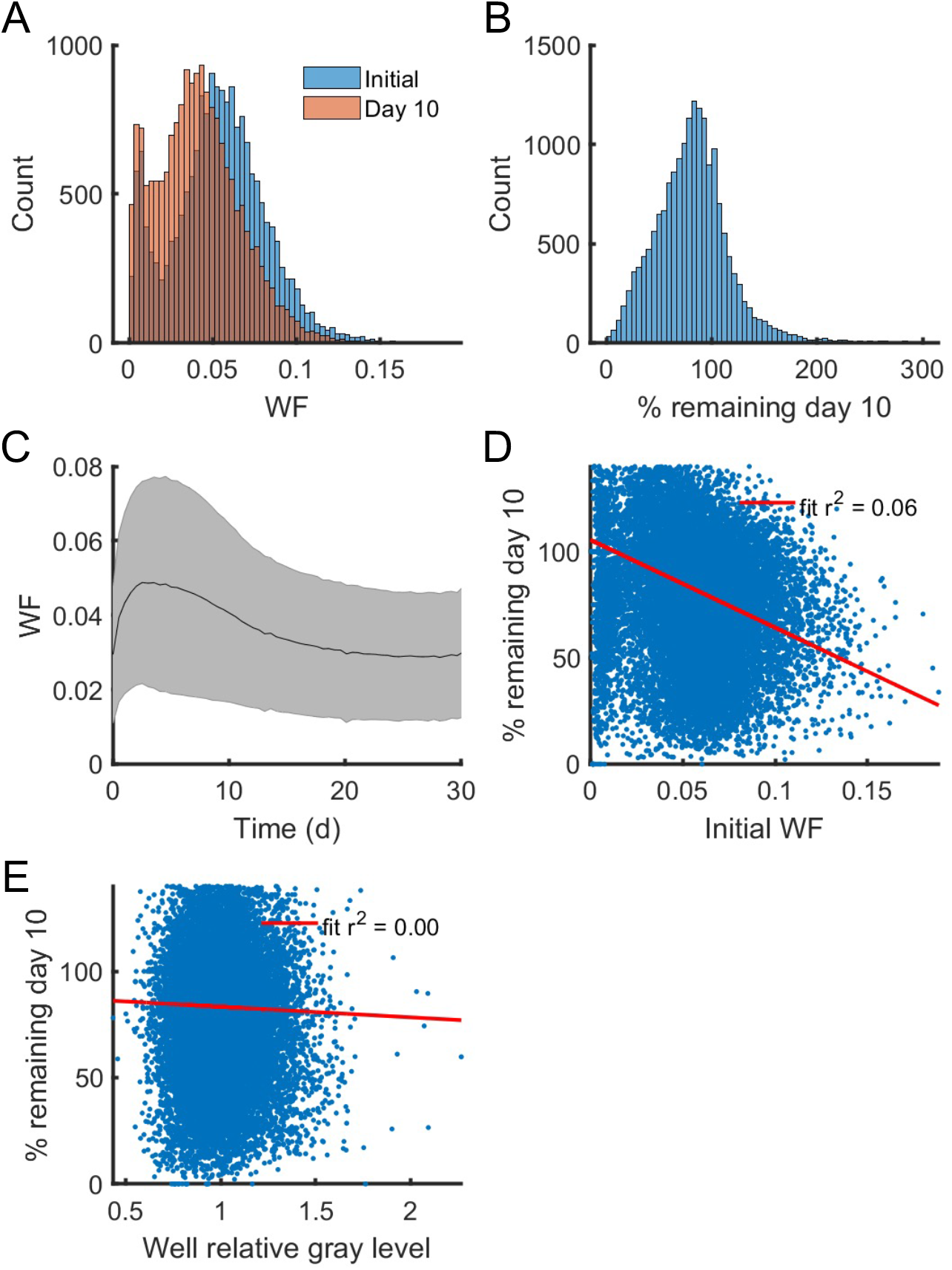
Quantification of worm escaping during screen experiments. (A) Histograms showing the distribution of worm fraction (WF) measurements initially (defined as the maximum value measured during the first 3 days) and at day 10. Day 10 mean is significantly lower (p < 0.001, paired t-test). (B) Distribution of the day 10 WF as a percentage of the initial WF. (C) Mean and standard deviation WF over time. (D-E) correlations between the percent remaining at day 10 and the initial WF or relative gray intensity of the well.

**Appendix I – Figure 5:**
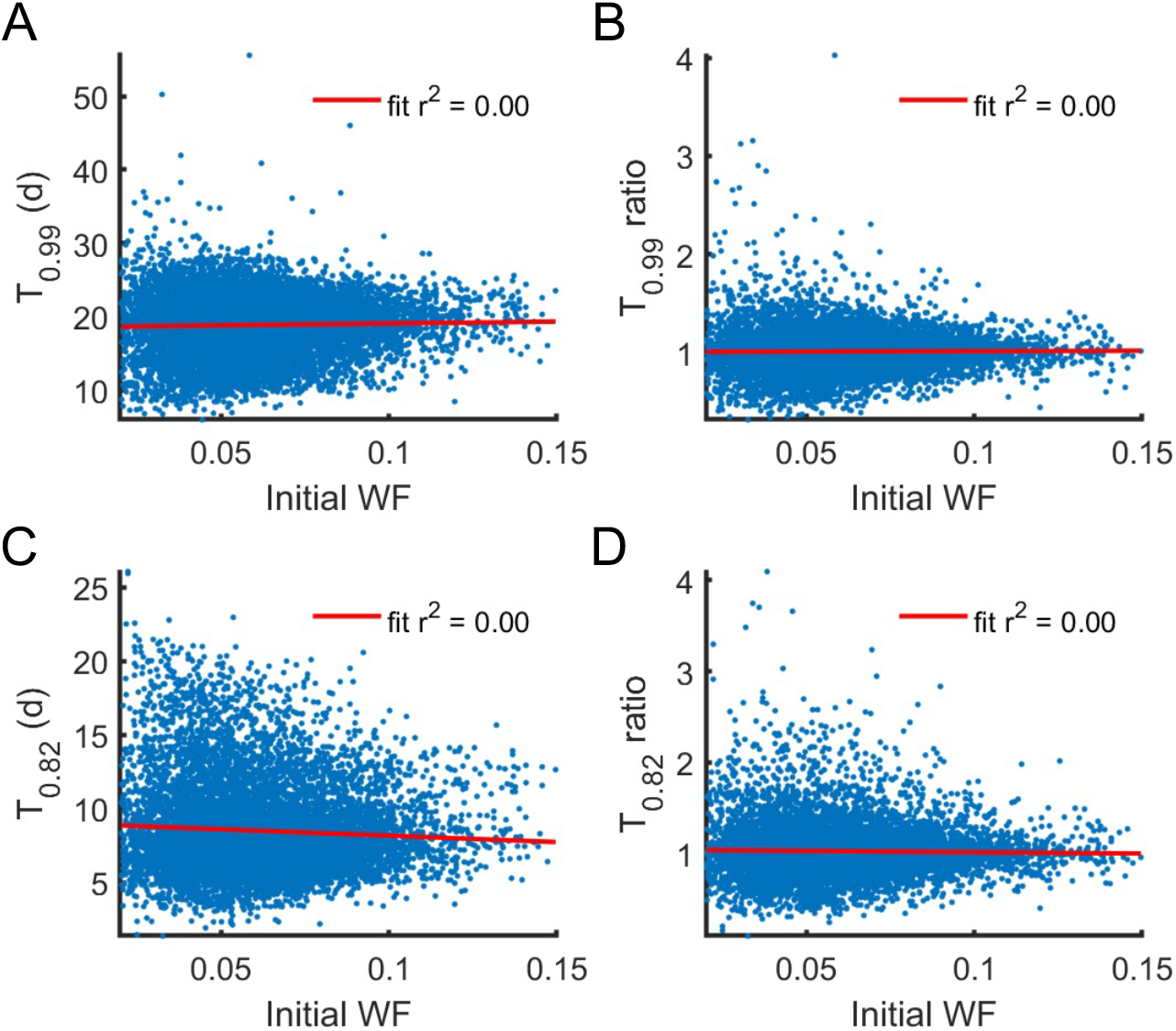
Initial worm density does not predict lifespan (A), lifespan ratio (B), healthspan (C), or healthspan ratio (D).

## Notes

### Competing Interest Statement

The authors have declared no competing interest.

## REFERENCES

Alisch, T., Crall, J.D., Kao, A.B., Zucker, D., and de Bivort, B.L. (2018). MAPLE (modular automated platform for large-scale experiments), a robot for integrated organism-handling and phenotyping. eLife 7, e37166.

Apfeld, J., and Kenyon, C. (1999). Regulation of lifespan by sensory perception in Caenorhabditis elegans. Nature 402, 804–809.

Bates, K., Le, K., and Lu, H. (2021). Deep learning for robust and flexible tracking in behavioral studies for C. elegans. bioRxiv, 2021.2002.2008.430359.

Bornhorst, J., Nustede, E.J., and Fudickar, S. (2019). Mass Surveilance of C. elegans— Smartphone-Based DIY Microscope and Machine-Learning-Based Approach for Worm Detection. Sensors 19.

Churgin, M.A., Jung, S.-K., Yu, C.-C., Chen, X., Raizen, D.M., and Fang-Yen, C. (2017). Longitudinal imaging of *Caenorhabditis elegans* in a microfabricated device reveals variation in behavioral decline during aging. eLife 6, e26652.

Cook, D.E., Zdraljevic, S., Roberts, J.P., and Andersen, E.C. (2017). CeNDR, the Caenorhabditis elegans natural diversity resource. Nucleic Acids Res 45, D650–D657.

Fraser, A.G., Kamath, R.S., Zipperlen, P., Martinez-Campos, M., Sohrmann, M., and Ahringer, J. (2000). Functional genomic analysis of C. elegans chromosome I by systematic RNA interference. Nature 408, 325–330.

Friedman, D.B., and Johnson, T.E. (1988). Three mutants that extend both mean and maximum life span of the nematode, *Caenorhabditis elegans*, define the age-1 gene. Journal of gerontology 43, B102–109.

Hakim, A., Mor, Y., Toker, I.A., Levine, A., Neuhof, M., Markovitz, Y., and Rechavi, O. (2018). WorMachine: machine learning-based phenotypic analysis tool for worms. BMC Biol 16, 8–8.

Hamilton, B., Dong, Y., Shindo, M., Liu, W., Odell, I., Ruvkun, G., and Lee, S.S. (2005). A systematic RNAi screen for longevity genes in C. elegans. Genes & Development 19, 1544–1555.

Hansen, M., Hsu, A.-L., Dillin, A., and Kenyon, C. (2005). New Genes Tied to Endocrine, Metabolic, and Dietary Regulation of Lifespan from a Caenorhabditis elegans Genomic RNAi Screen. PLOS Genetics 1, e17.

Herskind, A.M., McGue, M., Holm, N.V., Sørensen, T.I., Harvald, B., and Vaupel, J.W. (1996). The heritability of human longevity: a population-based study of 2872 Danish twin pairs born 1870-1900. Human genetics 97, 319–323.

Jagadeesan, S., and Hakkim, A. (2018). RNAi Screening: Automated High-Throughput Liquid RNAi Screening in Caenorhabditis elegans. Current protocols in molecular biology 124, e65.

Jushaj, A., Churgin, M., Yao, B., De La Torre, M., Fang-Yen, C., and Temmerman, L. (2020). Optimized criteria for locomotion-based healthspan evaluation in *C. elegans* using the WorMotel system. PLOS ONE 15, e0229583.

Kaeberlein, M. (2018). How healthy is the healthspan concept? GeroScience 40, 361–364.

Kamath, R.S., Fraser, A.G., Dong, Y., Poulin, G., Durbin, R., Gotta, M., Kanapin, A., Le Bot, N., Moreno, S., Sohrmann, M., et al. (2003). Systematic functional analysis of the *Caenorhabditis elegans* genome using RNAi. Nature 421, 231–237.

Kenyon, C., Chang, J., Gensch, E., Rudner, A., and Tabtiang, R. (1993). A *C. elegans* mutant that lives twice as long as wild type. Nature 366, 461–464.

Kenyon, C.J. (2010). The genetics of ageing. Nature 464, 504–512.

Lee, D., Zdraljevic, S., Stevens, L., Wang, Y., Tanny, R.E., Crombie, T.A., Cook, D.E., Webster, A.K., Chirakar, R., Baugh, L.R., et al. (2021). Balancing selection maintains hyper-divergent haplotypes in *Caenorhabditis elegans*. Nature Ecology & Evolution 5, 794–807.

Lee, S.S., Lee, R.Y.N., Fraser, A.G., Kamath, R.S., Ahringer, J., and Ruvkun, G. (2003). A systematic RNAi screen identifies a critical role for mitochondria in C. elegans longevity. Nature Genetics 33, 40–48.

Lucanic, M., Plummer, W.T., Chen, E., Harke, J., Foulger, A.C., Onken, B., Coleman-Hulbert, A.L., Dumas, K.J., Guo, S., Johnson, E., et al. (2017). Impact of genetic background and experimental reproducibility on identifying chemical compounds with robust longevity effects. Nature Communications 8, 14256.

Luyten, W., Antal, P., Braeckman, B.P., Bundy, J., Cirulli, F., Fang-Yen, C., Fuellen, G., Leroi, A., Liu, Q., Martorell, P., et al. (2016). Ageing with elegans: a research proposal to map healthspan pathways. Biogerontology 17, 771–782.

Mack, H.I.D., Heimbucher, T., and Murphy, C.T. (2018). The nematode Caenorhabditis elegans as a model for aging research. Drug Discovery Today: Disease Models 27, 3–13.

Mark, L., Monica, D., Patrick, P., Max, G., Jason, K., Carolina, I.-V., June, H., Shaunak, K., Elizabeth, C., Shobhna, P., et al. (2021). Protocol Exchange.

Olsen, A., Vantipalli, M.C., and Lithgow, G.J. (2006). Using Caenorhabditis elegans as a model for aging and age-related diseases. Ann N Y Acad Sci 1067, 120–128.

Paszke, A., Gross, S., Massa, F., Lerer, A., Bradbury, J., Chanan, G., Killeen, T., Lin, Z., Gimelshein, N., Antiga, L., et al. (2019). PyTorch: An Imperative Style, High-Performance Deep Learning Library. In NeurIPS.

Pitt, J.N., Strait, N.L., Vayndorf, E.M., Blue, B.W., Tran, C.H., Davis, B.E.M., Huang, K., Johnson, B.J., Lim, K.M., Liu, S., et al. (2019). WormBot, an open-source robotics platform for survival and behavior analysis in *C. elegans*. GeroScience 41, 961–973.

Pittman, W.E., Sinha, D.B., Zhang, W.B., Kinser, H.E., and Pincus, Z. (2017). A simple culture system for long-term imaging of individual *C. elegans*. Lab on a Chip 17, 3909–3920.

Rahman, M., Edwards, H., Birze, N., Gabrilska, R., Rumbaugh, K.P., Blawzdziewicz, J., Szewczyk, N.J., Driscoll, M., and Vanapalli, S.A. (2020). NemaLife chip: a micropillarbased microfluidic culture device optimized for aging studies in crawling C. elegans. Scientific Reports 10, 16190.

Saberi-Bosari, S., Huayta, J., and San-Miguel, A. (2018). A microfluidic platform for lifelong high-resolution and high throughput imaging of subtle aging phenotypes in C. elegans. Lab on a Chip 18, 3090–3100.

Stiernagle, T. (2006). Maintenance of *C. elegans*. In Wormbook, T.C.e.R. Community, ed.

Stroustrup, N., Ulmschneider, B.E., Nash, Z.M., López-Moyado, I.F., Apfeld, J., and Fontana, W. (2013). The *Caenorhabditis elegans* Lifespan Machine. Nature Methods 10, 665.

Uno, M., and Nishida, E. (2016). Lifespan-regulating genes in C. elegans. npj Aging and Mechanisms of Disease 2, 16010.

Xian, B., Shen, J., Chen, W., Sun, N., Qiao, N., Jiang, D., Yu, T., Men, Y., Han, Z., Pang, Y., et al. (2013). WormFarm: a quantitative control and measurement device toward automated Caenorhabditis elegans aging analysis. Aging Cell 12, 398–409.

Zhang, William B., Sinha, Drew B., Pittman, William E., Hvatum, E., Stroustrup, N., and Pincus, Z. (2016). Extended Twilight among Isogenic *C. elegans* Causes a Disproportionate Scaling between Lifespan and Health. Cell Systems 3, 333–345.e334.

